# Region-specific gene expression changes associated to oleoylethanolamide-induced attenuation of alcohol self-administration

**DOI:** 10.1101/2023.10.02.560440

**Authors:** Macarena González-Portilla, Sandra Montagud-Romero, Susana Mellado, Fernando Rodríguez de Fonseca, María Pascual, Marta Rodríguez-Arias

## Abstract

Oleoylethanolamide (OEA) is a lipid with anti-inflammatory activity that modulates multiple reward-related behaviors. Previous studies have shown that OEA treatment reduces alcohol self-administration (SA) while inhibiting alcohol-induced inflammatory signaling. Nevertheless, the specific mechanisms that OEA is targeting to achieve these effects have not been widely explored. Here, we tested the effects of OEA treatment during alcohol SA, extinction or previous to cue-induced reinstatement of alcohol seeking. In addition, we measured gene expression changes in the striatum and hippocampus of relevant receptors for alcohol consumption (DrD1, DrD2, CNR1, OPRM1) as well as immune-related proteins (IL-6, IL-1β, TLR4) and the brain-derived neurotrophic factor (BDNF). Our results confirmed that when administered contingently, systemic OEA administration reduced alcohol SA and attenuated cue-induced reinstatement. Interestingly, we also observed that OEA treatment reduced the number of sessions needed for extinction of alcohol seeking. Biochemical analyses showed region-specific OEA dopaminergic and cannabinoid receptor gene expression alterations. Also, OEA treatment modulated the long-term immune response and increased BDNF expression. These results suggest that boosting OEA levels may be an effective strategy for reducing alcohol SA and preventing relapse.

## 1. Introduction

Alcohol use disorder (AUD) is the most prevalent substance use disorder, causing an important impact on public health and society (Carvalho et al., 2019). Among the most frequent comorbidities of AUD are depression and anxiety disorders (Kathryn Mchugh & Weiss, 2019). The available pharmacological therapies have modest effects on reducing the risk of relapse and do not target the negative emotional state related to alcohol abstinence (Ray et al., 2018).

In recent years, lipid-signaling molecules have emerged as important modulators of reward-related behavior (Sagheddu et al., 2020). Oleoylethanolamide (OEA) is an endogenous molecule derived from oleic acid which functions as a satiety signal (Rodríguez De Fonseca et al., 2001; Schwartz et al., 2008). OEA has a common biosynthetic and degradation pathway as other N-acyl-ethanolamines (NAEs), anandamide (AEA) and palmitoylethanolamide (PEA), (Pagotto et al., 2006). All NAEs are synthesized on demand from lipid precursors and its deactivation depends mainly on hydrolysis by the enzyme fatty acid amide hydrolase (FAAH). OEA levels fluctuate during the day according mainly to nutritional status. In this regard, OEA levels increase after feeding, and especially upon intake of lipids (Schwartz et al., 2008). As a high-caloric substance, alcohol has also been proved to rapidly stimulate OEA production in enterocytes of the small intestine (Bilbao et al., 2016). Recently, a clinical study has proved that acylethanolamines levels, including OEA, are altered in AUD patients (Best et al., 2020; Garcia-Marchena et al., 2017).

In preclinical studies, OEA has been shown to affect the motivation for voluntary alcohol intake using animal models of both chronic and acute alcohol drinking. Using the self-administration (SA) and the two-bottle choice paradigms, OEA administration reduced alcohol drinking at 5, 10 and 20mg/kg doses (Bilbao et al., 2016; Sánchez-Marín et al., 2022). OEA also was effective in blocking cue-induced reinstatement of alcohol-seeking behavior (Bilbao et al., 2016). In addition, OEA has been proved to reduce the negative affect associated with alcohol withdrawal (Bilbao et al., 2016). These beneficial effects have been argued to depend on peripheral mechanisms since direct OEA infusion in the nucleus accumbens does not affect alcohol consumption. Also, sensory deafferentation on mice abolished the OEA-induced reduction in alcohol drinking (Bilbao et al., 2016). This body of research suggests that exogenous administration of OEA may act on peripheral receptors and signal to the central nervous system. In addition, OEA may be affecting alcohol-related behavior by its activity on the immune response. OEA interacts with peroxisome proliferator-activated receptors (PPAR- α) receptors expressed in immune cells and alters the transcription of target genes involved in the immune response (Guzman et al., 2004). Concerning alcohol, numerous studies have proved that OEA prevents inflammatory signaling induced by alcohol exposure (Orio et al., 2019).

To date, the effects of OEA on central gene expression associated to alcohol consumption have not been described. To further characterize the effects of OEA on voluntary alcohol drinking, we compared different OEA administration schedules at different timepoints during alcohol SA. Additionally, gene expression changes in the striatum and hippocampus of dopamine receptor (Dr)D1, DrD2, CNR1, OPRM, the immune-related genes, interleukin (IL)- 1β, IL-6, TLR4 and brain-derived neurotrophic factor (BDNF) were determined.

## 2. Methods and materials

### 2.1 Experimental design

OF1-strain male mice arrived at our laboratory postnatal day (PND) 21 (Charles River, France) and were housed in groups of four in standard plastic cages (27 x 27 x 14cm) under constant temperature and a reverse 12-h light/dark cycle (lights on at 7:00h). Mice were provided food and water *ad libitum* except during SA testing. All procedures were conducted in compliance with the guidelines of the European Council Directive 2010/63/EU regulating animal research and were approved by the local ethics committees of the University of Valencia (2019/VSC/PEA/0065). The experimental design is depicted in Figure 1.

**Figure 1.**
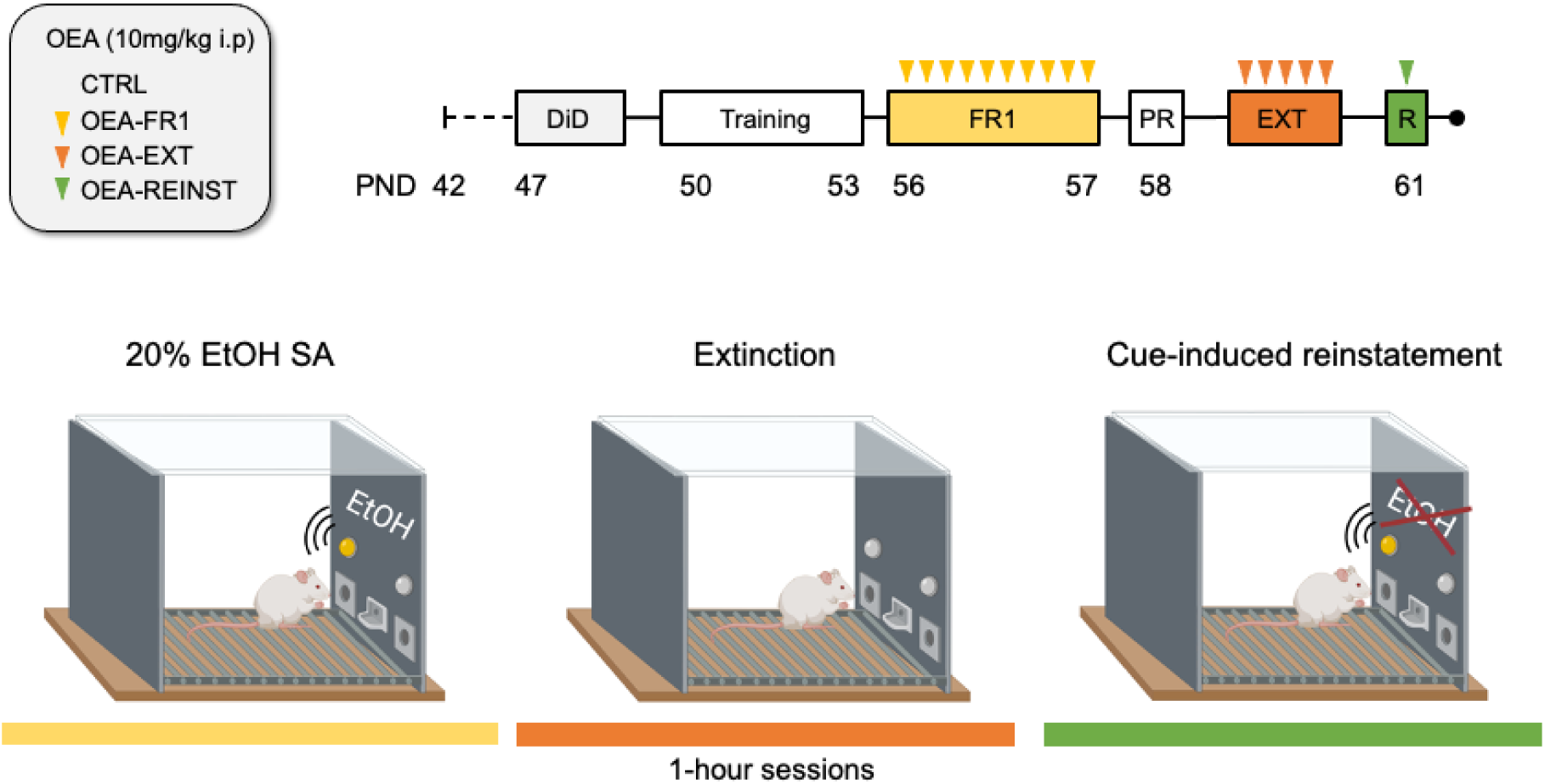
Experimental timeline. Mice were randomly assigned into four experimental groups according to OEA treatment (10mg/kg i.p): CTRL, OEA-FR1 (10 doses before each alcohol SA session), OEA-EXT (4-6 doses before each extinction session) and OEA-REINST (a single dose before the cue-induced reinstatement test). Mice were habituated to alcohol during a four-day Drinking in the dark procedure (DID). After SA training, mice self-administered 20% (v/v) alcohol during 10 days on a fixed-ratio 1 (FR1). When the period of extinction for alcohol seeking resumed, mice were tested on a single cue-induced reinstatement session in which alcohol solution was not delivered.

### 2.2 Drugs

OEA (10mg/kg, i.p.; synthesized as described in Rodríguez De Fonseca et al., (2001)) was dissolved in 5% Tween 80 in saline and injected 10 minutes before the corresponding timepoint according to the experimental condition. The doses were chosen according to previous studies in mice reporting effective therapeutic and anti-inflammatory effects (González-Portilla et al., 2023; Moya et al., 2021; Rivera et al., 2018; Sayd et al., 2015).

### 2.3 Habituation to alcohol

The Drinking in the Dark (DiD) paradigm was employed as a pre-exposure procedure to habituate mice to ethanol before starting the oral SA. Based on the basic paradigm of Rhodes et al. (2005), this voluntary drinking protocol consisted of two phases. On the first day mice were moved from their homecage for two hours to habituate to the individual cages and the drinking tubes. In the second phase of the protocol, mice received 3 days of 2h/day access to 20% (v/v) ethanol solution starting 3h after lights off. On day 4, the procedure extended for 4 h. After the each DiD session, mice returned to their homecage.

### 2.4 Oral alcohol self-administration (SA)

This procedure is based on that employed by Navarrete et al., (2014). Voluntary oral alcohol SA administration was carried out in eight modular operant chambers (MED Associated Inc., Georgia, VT, USA) equipped with a chamber light, two nose-poke holes, one fluid receptacle, one syringe pump, one stimulus light and one buzzer. The chambers were placed inside sound-attenuated cubicles. Designated active nose-pokes delivered 36μl of fluid associated to a 0.5s light cue and a 0.5s buzzer beep, which was followed by a 6s time-out period. Inactive nose-pokes triggered no event. Software package (Cibertec, SA, Spain) controlled stimulus and fluid delivery and recorded operant responses.

The procedure consisted of five phases: training, fixed ratio 1 (FR1), progressive ratio (PR) 20% (v/v) ethanol solution, extinction, and reinstatement. Alcohol SA was assessed by daily operant sessions of 60-minute duration.

#### Training (15 days)

Mice were trained to poke to the active hole for 20% (v/v) ethanol solution delivery (20 μl) under a FR1. Inclusion criteria for the next step in the procedure comprised 60 % discrimination for active, over inactive nose-poke responding across the three last days of this phase. Mice that met these criteria were randomly divided into four experimental groups: CTRL, OEA-FR1, OEA-EXT and OEA-REINST groups (n=10/12).

#### FR1 (10 days)

The number of effective responses and 20% ethanol (v/v) consumption (μl) were measured under a FR1 for 10 daily sessions. After each session, the alcohol solution that remained in the receptacle was collected and measured with a micropipette.

#### PR (1 day)

Mice undertook a single 2h long PR session in which the response requirement necessary to obtain one reinforcement escalated according to the following series: 1-2-3-5-12-18-27-40-60- 90-135-200-300-450-675-1000. Breaking point (BP) was defined as the highest number of nose-pokes each mouse performed to earn one reinforcement. This value was used to quantify motivational strength.

#### Extinction sessions

All mice progressed to the extinction phase, which consisted of removal of the cue light, buzzer, and the reward (alcohol) delivery after a response on the active nose-poke during the 60- minute session.

The extinction phase continued until the average in each experimental group reached the criterion (at least 50% decrease in active nose poke responses for at least three consecutive days).

#### Cue-induced reinstatement of alcohol seeking (1 day)

Following the extinction phase, a 60-minute single session of cue-induced reinstatement was performed. Effective responses on the active nose-poke were followed by presentation of cue light and buzzer stimuli in the absence of alcohol delivery. For this session, fluid receptable was primed with alcohol solution.

### 2.5 Tissue sampling and biochemical analyses

Mice were sacrificed by cervical dislocation. The striatum and hippocampus were precisely dissected out using a coronal brain matrix. Tissue samples were stored at -80°C until the qRT-PCR assay was performed.

#### 2.5.1 RNA Isolation, Reverse Transcription, and Quantitative RT-PCR

Striatum and hippocampus were lysed in Tri-Reagent solution (Sigma-Aldrich, St. Louis, USA) and total RNA was isolated according to the manufacturer’s instructions. Then, the mRNA was reverse-transcribed by the High Capacity cDNA Reverse Transcription Kit (Applied Biosystems, Massachusetts, USA). Amplification of the target and housekeeping genes was completed employing the AceQ ® qPCR SYBR Green Master Mix (NeoBiotech, Nanterre, France) and TaqManTM Fast Advanced Master Mix (Applied Biosystems, Massachusetts, USA), following the manufacturer’s instructions in a QuantStudioTM 5 Real-Time PCR System (Applied Biosystems, Massachusetts, USA). The mRNA level of housekeeping genes (cyclophilin A and β-glucuronidase) was used as an internal control for the normalization of the analyzed genes. All the RT-qPCR runs included non-template controls (NTCs). Experiments were performed in triplicates. Quantification of expression (fold change) from the Cq data was calculated by the QuanStudioTM Design & Analysis Software (Applied Biosystems). Details of the nucleotide sequences and the assay codes of the used primers are detailed in the Supplementary Material (Tables S1 and S2).

### 2.6 Statistical analyses

Data relating to body weight were analyzed by a mixed ANOVA with one between-subjects variable “Treatment” with 4 levels (CTRL, OEA-FR1, OEA-EXT, OEA-REINST) and a within variable “Weeks” with 9 levels (1-9).

For the SA, alcohol consumption and active nose poke during responses during FR1 were analyzed individually by a two-way ANOVA with one between-subject’s variable –Treatment (CTRL, OEA) and a within-subject’s variable –Days, with ten levels of FR1 schedule.

A one-way ANOVA with a between-subject’s variable – Treatment (CTRL, OEA-FR1, OEA-EXT and OEA-REINST) – was employed to analyze alcohol consumption and breaking point values during the PR session.

The time required for meeting the extinction criteria in each animal was analyzed by means of the Kaplan–Meier test with Breslow (generalized Wilcoxon) comparisons. When a mouse did not meet the extinction criteria during the extinction period, we assigned the number of days required for extinction for the group as a whole.

To test for the cue-induced reinstatement, the number of active nose poke responses were analyzed with a two-way ANOVA with one between-subjects’ variable – Treatment, with two levels (CTRL, OEA)– and one within subjects’ variable with two levels (Day: extinction and cue-induced reinstatement).

The gene expression data were analyzed by a one-way ANOVA with one between variables, Treatment, with four levels (CTRL, OEA-FR1, OEA-EXT, OEA-REINST). Bonferroni post-hoc tests were also analyzed. Statistical analyses were performed using SPSS Statistics v23.

## 3. Results

### 3.1 Bodyweight

Given the reported effects of OEA on feeding behavior, we obtained a weekly measure of mice bodyweight. The ANOVA revealed a significant effect of the variable week [F(8,336)=98.819, (p<0.001)], as mice exhibited an increased bodyweight across weeks compared to the first measurement (p<0.001), (Figure 2). No differences were found between groups.

**Figure 2.**
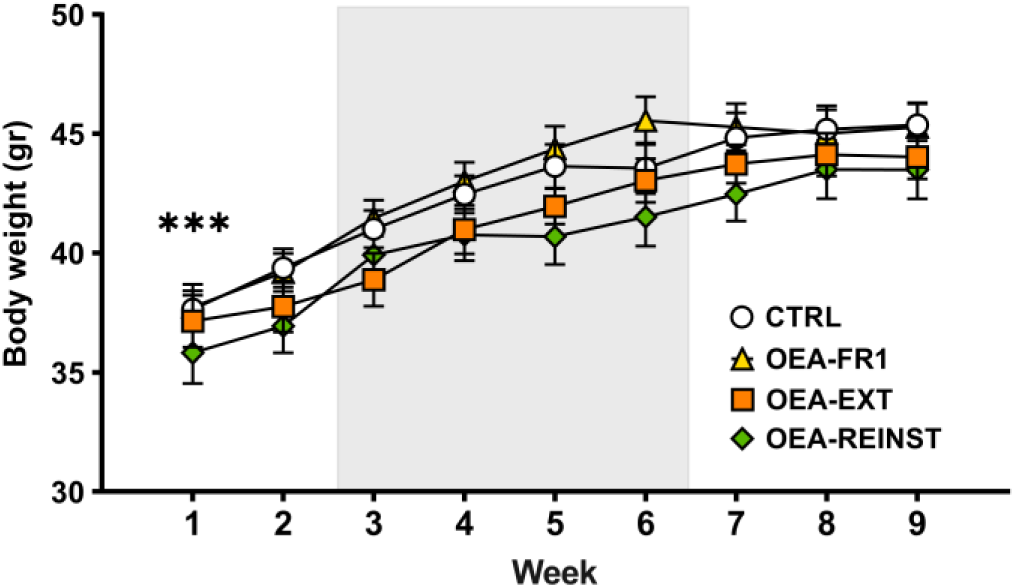
Body weight across the experiment. Grey area represents alcohol 20% self-administration period (training and fixed-ratio 1). Data are represented as the mean (±SEM) bodyweight measured weekly. *** p < 0.001 significant difference with respect to Week 1.

### 3.2 Oral alcohol SA

The ANOVA for the ethanol consumption during the FR1 schedule revealed a significant effect of the variable Treatment [F(1,44) = 4,091; p = 0.049]. A lower EtOH consumption was observed in OEA-FR1 mice compared to the CTRL, OEA-EXT and OEA-REINST mice (p = 0.049), (Figure 3).

**Figure 3.**
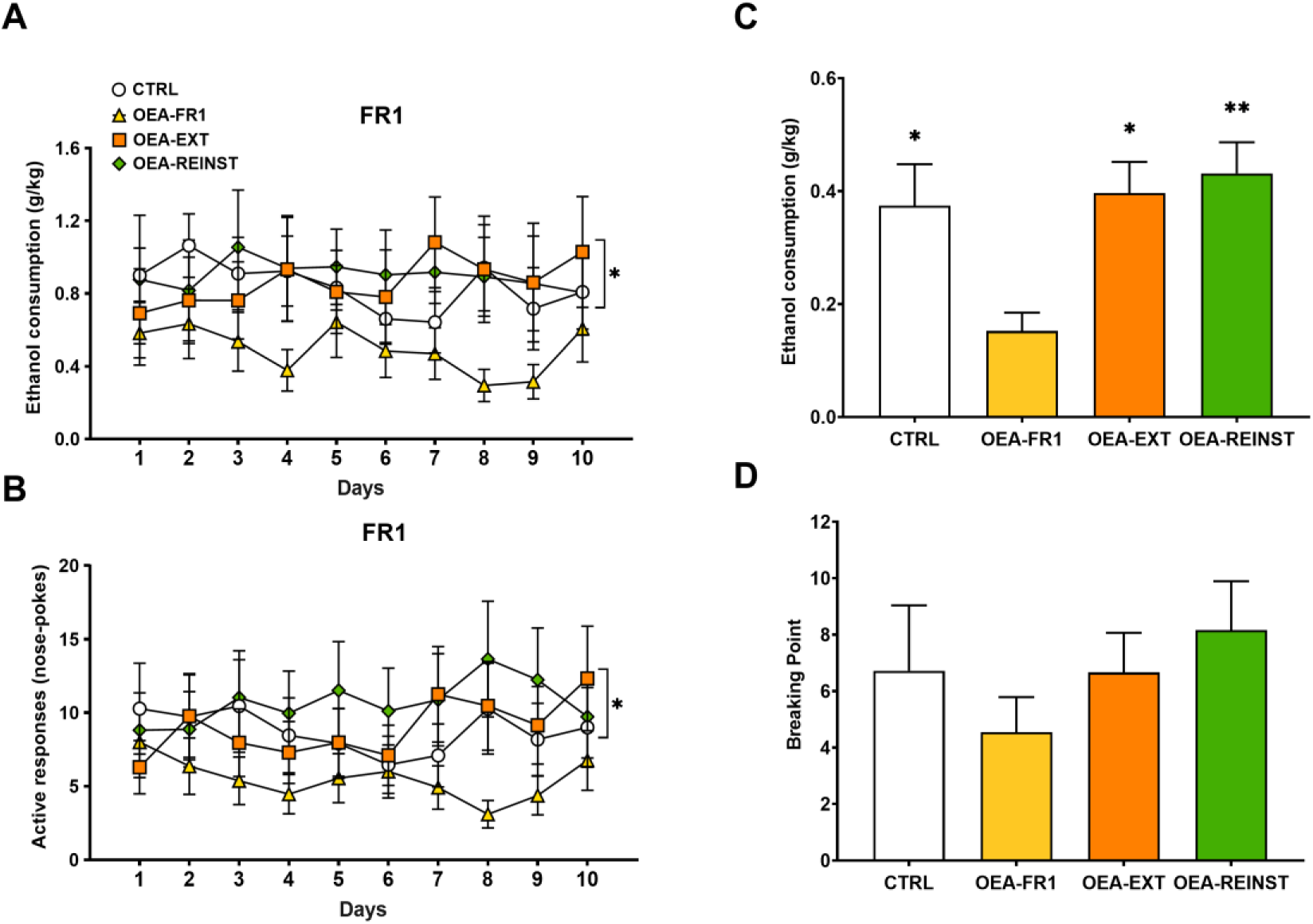
Effects of OEA treatment on oral alcohol self-administration. The dots represent means and the vertical lines ± SEM of (a) 20% EtOH consumption during the FR1 period, (b) active nose poke responses during FR1 period. (c) EtOH consumption during the PR session, and (d) breaking point values during the PR session. * p < 0.05, ** p < 0.01 significant difference with respect to the OEA-FR1 group.

Regarding active responses, the ANOVA showed an effect of the variable Treatment [F(1,44) = 4,827; p = 0.033]. Mice in the OEA-FR1 group performed fewer active responses compared to the CTRL, OEA-EXT and OEA-REINST mice (p = 0.033).

During the PR, the ANOVA revealed a significant effect of the variable “Treatment” [F(3, 42)=57.957 (p<0.001). Post-hoc comparisons showed that alcohol consumption was decreased in OEA-FR1 mice compared to CTRL (p=0.52), OEA-EXT (p=0.021) and OEA-REINST (p=0.006) groups.

With regards to the time required to extinguish alcohol SA, the OEA-EXT group required a mean number of 3.4 sessions, while the CTRL, OEA-FR1 and OEA-REINST groups required a mean number of 4.8, 5,5 and 4.9 sessions, respectively (Figure 4A). The Kaplan-Meier analysis revealed that the OEA-EXT group required significantly less sessions than the CTRL (χ2 = 4.322; p = 0.038), OEA-FR1 (χ2 = 9.549; p = 0.002) and OEA-REINST (χ2 = 4.933; p = 0.026) to meet the extinction criteria.

**Figure 4.**
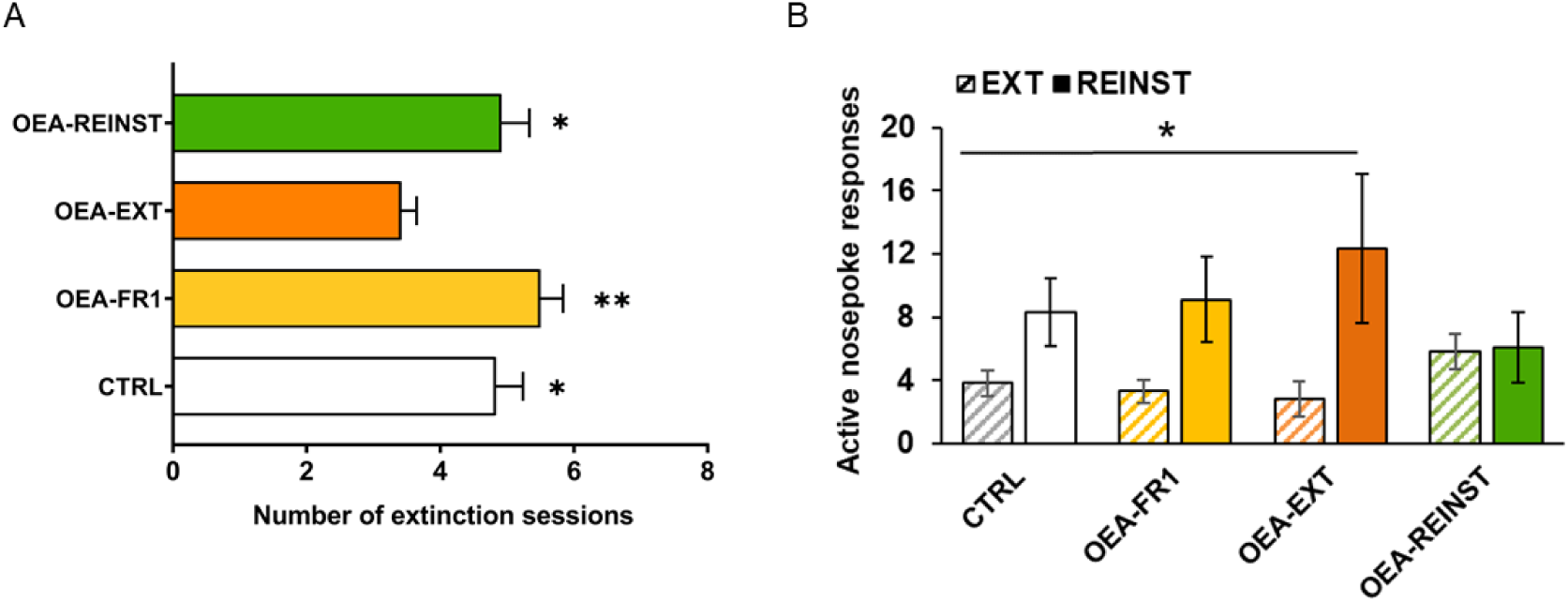
(a) Extinction. The bars represent the total value (±S.E.M) of the number of sessions required for alcohol drug-seeking to be extinguished. * p < 0.05, * p < 0.01significant difference with respect to the OEA-EXT group. (b) Effects of OEA treatment on cue-induced reinstatement of alcohol-seeking. Mean active nose pokes responses during the last three extinction sessions and on the cue-induced reinstatement test. The columns represent means and the vertical lines ± SEM of active nosepoke responses. * p < 0.05, significant difference with respect to EXT.

Finally, we tested the effects of OEA on drug-seeking behavior induced by contextual cues. The ANOVA revealed an effect of the variable Day [F(1,43) = 4,825; p = 0.033] and the interaction Day x Treatment [F(1,43) = 4,160; p = 0.048], (Figure 4B). The active responses induced by the alcohol-associated stimuli on the reinstatement test were significantly increased with respect to the last three days of extinction. The CTRL, OEA-FR1 and OEA-EXT groups exhibited increased active responses with respect to the extinction period (p<0.001). No differences in the number of active responses were observed for mice in the OEA-REINST group.

### 3.3 OEA affects gene expression in the striatum and hippocampus

The ANOVA revealed an effect of the variable Treatment for striatal DrD1 [F(3,28) = 4,595; p = 0.01], cannabinoid receptor-1 gene (CNR1) [F(3,28) = 3,539; p = 0.027] and OPRM1 [F(3,28) = 3,352; p = 0.033] gene expression. OEA administration in the OEA-FR1 group significantly decreased DrD1 mRNA expression compared to the CTRL (p = 0.01) and the OEA-REINST group (p = 0.052). Similarly, a decrease in CNR1 mRNA expression was observed in the striatum of OEA-FR1 mice compared to OEA-REINST mice (p = 0.024). In addition, we found increased levels of OPRM1 gene expression in the striatum in OEA-EXT mice compared to CTRL mice (p = 0.037), (Figure 5A).

**Figure 5.**
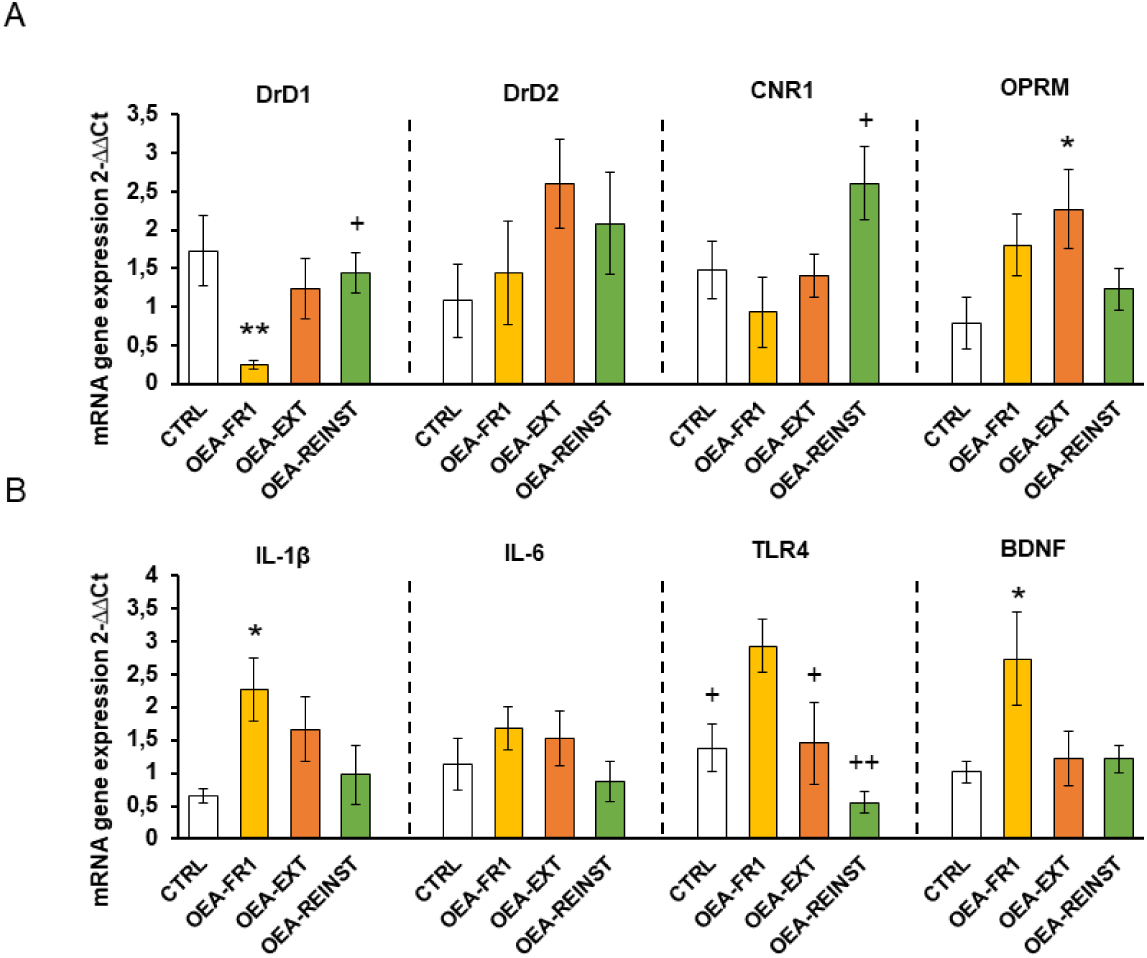
Effects of OEA administration on striatal gene expression of a) relevant receptors involved in alcohol reinstatement (DrD1, DrD2, CNR1 and OPRM) and (b) inflammatory genes, including IL-1β, IL-6 and TLR4, as well as the neurotrophic factor BDNF after alcohol cue-induced reinstatement. * p < 0.05 significant difference with respect to the CTRL group; + p < 0.05, ++ p < 0.01 significant difference with respect to the OEA-FR1 group.

Regarding the expression of inflammatory genes in the striatum, the ANOVA revealed an effect of the variable Treatment for IL-1β [F(3,28) = 3,67; p = 0.024], and TLR4 [F(3,28) = 6,873; p = 0.001]. We found increased mRNA expression of IL-1β in the striatum of OEA-FR1 mice compared to CTRL mice (p = 0.031). Similarly, gene expression of TLR4 was increased in the striatum in OEA-FR1 mice compared to CTRL (p = 0.42), OEA-EXT (p = 0.59) and OEA-REINST mice (p < 0.001), (Figure 5B).

Finally, the ANOVA for the striatal BDNF gene expression showed an effect of the variable Treatment [F(3,27) = 3,922; p = 0.019]. Gene expression analyses showed that OEA-FR1 mice exhibited increased BDNF mRNA levels compared to the CTRL mice (p < 0.30), (Figure1B).

The ANOVA revealed an effect of the variable Treatment for hippocampal DrD1 [F(3,28) = 5,341; p = 0.005], CNR1 [F(3,27) = 12,892; p < 0.001] and OPRM1 gene expression [F(3,28) = 4,514; p = 0.011]. An increase in DrD1 gene expression was observed in the hippocampus of OEA-FR1 compared to OEA-EXT mice (p = 0.004). Also, an increase in hippocampal CNR1 gene expression was observed in OEA-FR1 compared to the CTRL (p = 0.002), OEA-EXT (p < 0.001) and OEA-REINST groups (p = 0.10). In addition, OPRM1 gene expression was decreased in the hippocampus of OEA-FR1 mice compared to CTRL (p = 0.40), OEA-EXT (p=0.45) and OEA-REINST groups (p = 0.22), (Figure 6A).

**Figure 6.**
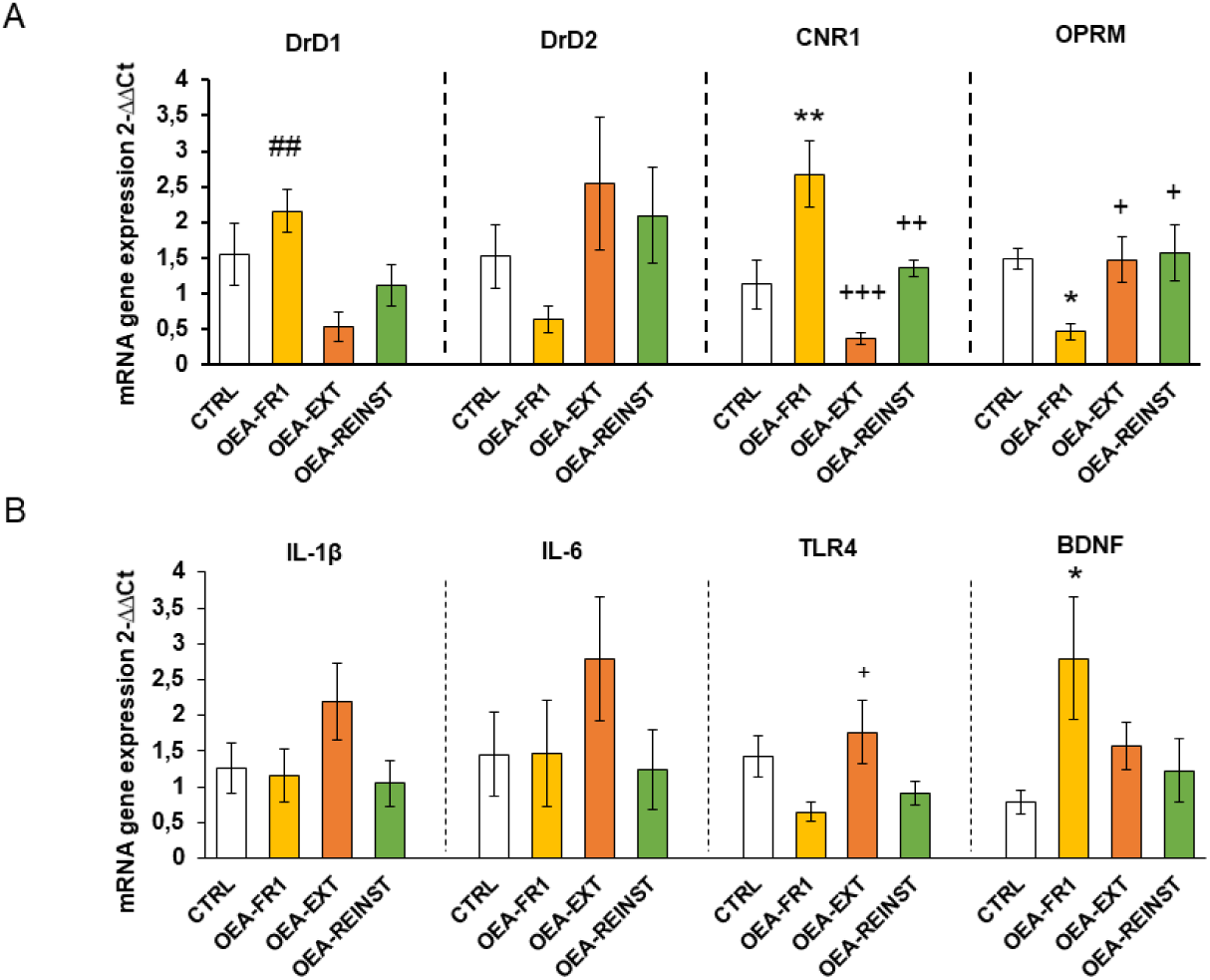
Effects of OEA administration on hippocampal gene expression of a) relevant receptors involved in alcohol reinstatement (DrD1, DrD2, CNR1 and OPRM) and (b) inflammatory genes, including IL-1β, IL-6 and TLR4, as well as the neurotrophic factor BDNF after alcohol cue-induced reinstatement. ** p < 0.01 significant differences with respect to the CTRL group; ## p < 0.01 significant difference with respect to the OEA-EXT group; +++ p < 0.001, ++ p < 0.01 significant difference with respect to the OEA-FR1 group.

Regarding the gene expression of inflammatory immune receptor, TLR4, in the hippocampus, the ANOVA revealed an effect of the variable Treatment [F(3,28) = 3,884; p = 0.019]. The results also showed a downregulation of its mRNA levels in OEA-FR1 mice compared to OEA-EXT mice, (Figure 6B).

Finally, the ANOVA for the hippocampal BDNF gene expression showed an effect of the variable Treatment [F(3,27) = 3,217; p = 0.38]. These analyses showed that OEA-FR1 mice exhibited increased BDNF mRNA levels compared to the CTRL mice (p = 0.38)

## 4. Discussion

Previous literature presents OEA as a lipid that modulates the response to natural rewards, such as food, as well as to drugs of abuse. In this study, we tested the attenuating effects of OEA on alcohol self-administration. We confirmed that acute OEA administration (10mg/kg) reduces alcohol consumption. We also observed that OEA treatment accelerates the extinction of alcohol-seeking behavior. Finally, we show that a single OEA dose blocks cue-induced reinstatement of alcohol seeking. In addition, the current study shed some light on the effects of OEA on striatal and hippocampal levels of relevant genes for alcohol-related behavior (DrD1, DrD2, CNR1, OPRM), neuroinflammatory processes (IL-1β, IL-6 and TLR4) and BDNF.

As previously reported, we observed that acute OEA administration before each session reduces operant responses for alcohol SA and consumption (Bilbao et al., 2016). Similar to the FR1 period, mice in the OEA-FR1 group exhibited decreased alcohol consumption during the PR session. Although not significant, a similar trend was observed in the breaking point values, which suggest a reduction in the motivation to obtain alcohol.

One of the most interesting findings of this study was the effect of OEA on the extinction of alcohol-associated memories. Given the evidence showing that OEA reduces the severity of alcohol withdrawal negative symptoms (Bilbao et al., 2016, Anton et al., 2016), we tested the effects of OEA during the extinction period (OEA-EXT group), in which neither alcohol nor alcohol-associated cues were available following an active response. Interestingly, we observed that OEA treatment before each extinction session shortened the time needed to extinguish alcohol-seeking behavior. To our knowledge, this is the first study to demonstrate an attenuating effect of OEA on extinction of alcohol-seeking behavior. These findings are in line with human studies showing that serum OEA levels are increased in AUD patients during alcohol withdrawal (García-Marchena et al., 2017). As a molecule with antidepressant properties, it is possible that OEA buffers the negative effects of alcohol abstinence on mood during the extinction period. Otherwise, OEA may also be exerting a cognitive enhancer effect that potentiate the key learning mechanisms that mediate extinction learning, thus facilitating the creation of new associations. A previous study reported an improvement in cognitive functions in a mice model of Wernicke-Korsakoff syndrome (Moya et al., 2021). In this sense, OEA may be accelerating the consolidation of the extinction learning while reducing the aversive symptomatology associated to alcohol deprivation, thus curbing alcohol-related motivational memories.

Consistent with a previous study, we also observed that a single dose was effective in blocking cue-induced reinstatement of alcohol seeking (Bilbao et al., 2016). The mechanisms through which OEA exerts an attenuating effect on cue-induced reinstatement may be diverse. It is possible that OEA improves the reward deficit induced by alcohol SA and ameliorates craving and depressive-like behavior which prevented reinstatement. To gain better insight into the neurotransmission system involved in OEA effects on alcohol-related behaviors we evaluated the transcriptional regulation in two brain regions of the reward system, the striatum and hippocampus.

In humans, postmortem studies show that the number of binding sites of dopamine DrD1 and CNR1 are increased in the striatum of individuals with a history of AUD (Hirth et al., 2016) Within the striatum, the receptor DrD1 is an important receptor is involved in dopamine signaling during alcohol craving and relapse. Our results showed that DrD1 gene expression was decreased in the striatum of OEA-FR1 mice. Consistent with these results, in a previous study we demonstrated that striatal DrD1 gene expression was decreased in mice that received OEA during each session of cocaine-induced conditioned place preference compared to the vehicle-treated mice (González-Portilla et al., 2022). It is important to remark that at the point of tissue collection, the time from the OEA treatment as well as the number of doses between groups differed. Likely, changes in dopaminergic receptors were not detected in the OEA-EXT group because mice received fewer doses of OEA (4-6). Concerning mice in the OEA-REINST group, we consider that a single dose of OEA may not impact DrD1/2 gene expression significantly. Taken together, our results show that OEA altered DrD1 gene expression in the striatum of mice that received OEA contingently to alcohol SA, which may be related to the reduction in alcohol consumption.

It has been demonstrated that the dopaminergic system is capable to interact with cannabinoid signaling, modulating inputs to the striatum (Peters et al., 2021). In this sense, it is possible that OEA reduces alcohol consumption by acting on these two neurotransmission systems in reward-related areas. In the striatum, decreased CNR1 gene expression was detected in OEA-FR1 mice only with respect to the OEA-REINST group. At 24hr after injection, OEA possibly has an opposite effect, increasing CNR1 gene expression due to competition for the catalytic function of FAAH (Ho et al., 2008). Multiple studies have consistently shown that genetic, or pharmacological blockade of CB1R reduces most alcohol-related behaviors such as alcohol drinking and reinstatement in the operant paradigm (de Bruin et al., 2011; Economidou et al., 2006; Hungud et al., 2003; Rodríguez De Fonseca et al., 1999). On the contrary, stimulating CB1R using cannabinoids or blocking the FAAH enzyme increases alcohol seeking and drinking (Cinar et al., 2020; Zhou et al., 2017). According to these results, we suggest that multiple doses in OEA-FR1 mice affects the dopaminergic and endocannabinoid system, diminishing the rewarding properties of alcohol and resulting in decreased alcohol intake.

We observed opposite changes to those of the striatum for DrD1 and CNR1 gene expression in the hippocampus. The molecular changes involving the dopaminergic and endocannabinoid systems, including downregulation of CB1 receptor, are complex and frequently region-specific (Lazenka et al., 2013; Soria-Gomez et al., 2021). Also, lipidomic analyses in alcohol self-administered mice show heterogeneous lipid profiles, including OEA, in multiple brain areas (Shahen-Zoabi et al., 2023).

Due to the implication of the opioid system in alcohol reward, we also examined OPRM1 gene expression. In the striatum, OEA-EXT group showed increased gene expression of OPRM1 compared to the CTRL group. The specific cause for this upregulation is unclear, although it may be attributable to the blunting effect of OEA on the negative emotional state resulting from alcohol abstinence, which affects opioid signaling (Hansson et al., 2019). Differently from the striatum, OEA-FR1 mice exhibited a downregulation of OPRM1 in the hippocampus. In fact, several studies have proved that mice lacking the OPRM1 gene exhibit a lower preference for alcohol and reduced self-administration compared to wild-type mice (Hall et al., 2001).

Another potential mechanism by which OEA may be exerting its effects is by modulating the alcohol-induced immune signaling. Multiple studies proved that OEA reduces the release of proinflammatory factors in different animal models of disease and after an immune challenging event (Comerota et al., 2023; Rivera et al., 2019). For example, exogenous OEA reduced the proinflammatory mediators IL-1β, IL-6, after an LPS injection (Sayd et al., 2015). An important signaling cascade of alcohol-induced immune response are TLR4 receptors (Pascual et al., Crews et al., 2017). Recent reports have proved the protective effects of OEA on acute and chronic activation of TLR4 by drug exposure (González-Portilla et al., 2023; Moya et al., 2021). Up to date, most studies have measured protein levels of numerous cytokines and chemokines, showing that OEA treatment counteracts the alcohol-induced increase in these immune factors (Orio et al., 2019). In our study, we assessed transcription changes in several inflammatory-related genes two weeks after alcohol SA resumed. In this sense, we aim to detect lasting immune signatures associated to alcohol abstinence and craving rather than an acute alcohol-induced inflammatory response. In the striatum, we observed an upregulation of immune genes (e.g., IL-1β and TLR4) in the OEA-FR1 mice, compared to the CTRL group. In the hippocampus, the upregulation of TLR4 expression was specific to the OEA-EXT group compared to the OEA-FR1 mice. Since the peak inflammatory signaling occurred concomitantly to the alcohol consumption, we presume that during the FR1 period, the immune response was attenuated in the OEA-FR1 mice by the OEA anti-inflammatory activity. In this sense, it is possible that the detected increase in TLR4 results from a rebound effect. Given the timepoint when the neuroimmune gene transcript was obtained this increase in the OEA-FR1 group likely reflect homeostatic interactions between different immune systems.

Finally, we also examined the neuroprotective growth factor BDNF which is an important regulator of activity-driven synaptic plasticity (Lu et al., 2003). The highest number of BDNF receptor is found in the hippocampus (Masana et al., 1993). Previous studies show that OEA treatment restores BDNF following cerebral ischemia (Yang et al., 2015) or after stress exposure (Jin et al., 2015). Alcohol exposure affects BDNF gene expression in a time and dose-dependent manner. In our study, we observed that all OEA-FR1 mice exhibited higher levels of BDNF gene expression compared to the CTRL group in the striatum and hippocampus. It is worth noting that the highest gene expression of BDNF was observed in the hippocampus of OEA-FR1 mice which is the group that received the highest number of doses of OEA (10 daily i.p injections). These results are consistent with a previous study that reported enhanced BDNF gene expression in the hippocampus of OEA-treated mice exposed to ethanol binge drinking compared to the vehicle-treated mice (Silva-Peña et al., 2019). In this sense, it is possible that the administration of multiple doses of exogenous OEA has a buffering effect on the alcohol-induced neuronal toxicity by enhancing BDNF expression.

All in all, we propose that the observed gene expression alterations are associated to the oleoylethanolamide-induced reduction in alcohol SA, extinction of alcohol-seeking and attenuated alcohol reinstatement. Nevertheless, different central and peripheral mechanisms may be contributing to this effect. Additionally, it is important to remark that many of the changes in dopaminergic, cannabinoid receptor and immune-related gene expression may be secondary to the OEA-induced reduction of alcohol consumption in the OEA-FR1 group.

## 5. Conclusions

The present findings support the role of OEA as a homeostatic mediator modulating multiple alcohol-related behaviors. Taken together, OEA administration reduced alcohol SA, accelerated extinction of alcohol-seeking and attenuated cue-induced reinstatement. The results of the present study and our previous work indicate that OEA is the most effective in decreasing drug reward when administered contingently. Biochemical analyses suggest that OEA administration affects gene expression of dopaminergic, cannabinoid, opioid and immune-related factors in the striatum and hippocampus, key brain areas for reward processing. The impact of OEA on gene expression was heterogeneous, suggesting that both the quantity of doses and the timing of administration are crucial factors. Given these results, further studies should aim to elucidate the specific mechanisms OEA is targeting to modulate alcohol SA.

## Supplementary Material

**Table S1.**
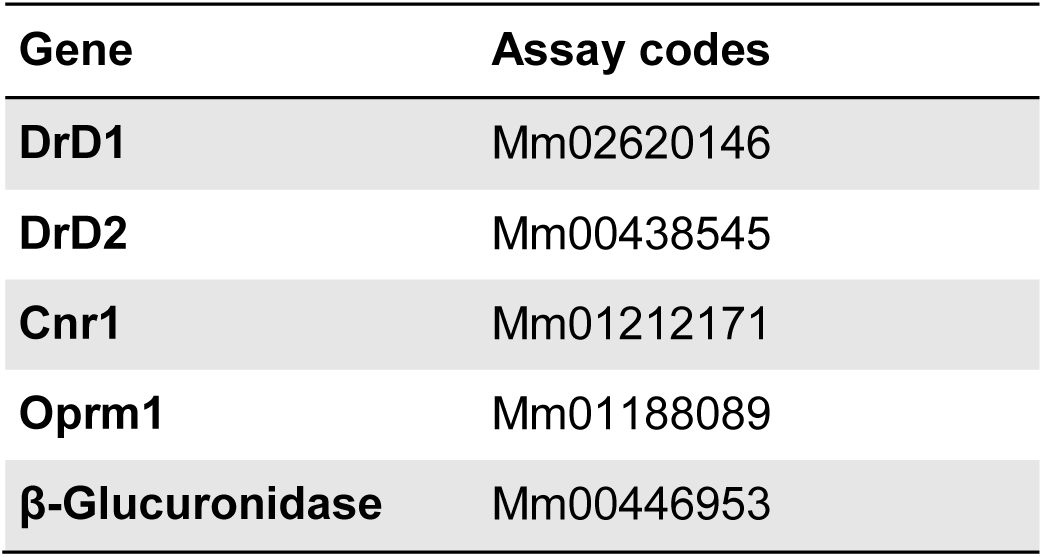
Assay codes of the primers used for the RT-PCR of genes.

**Table S1.**
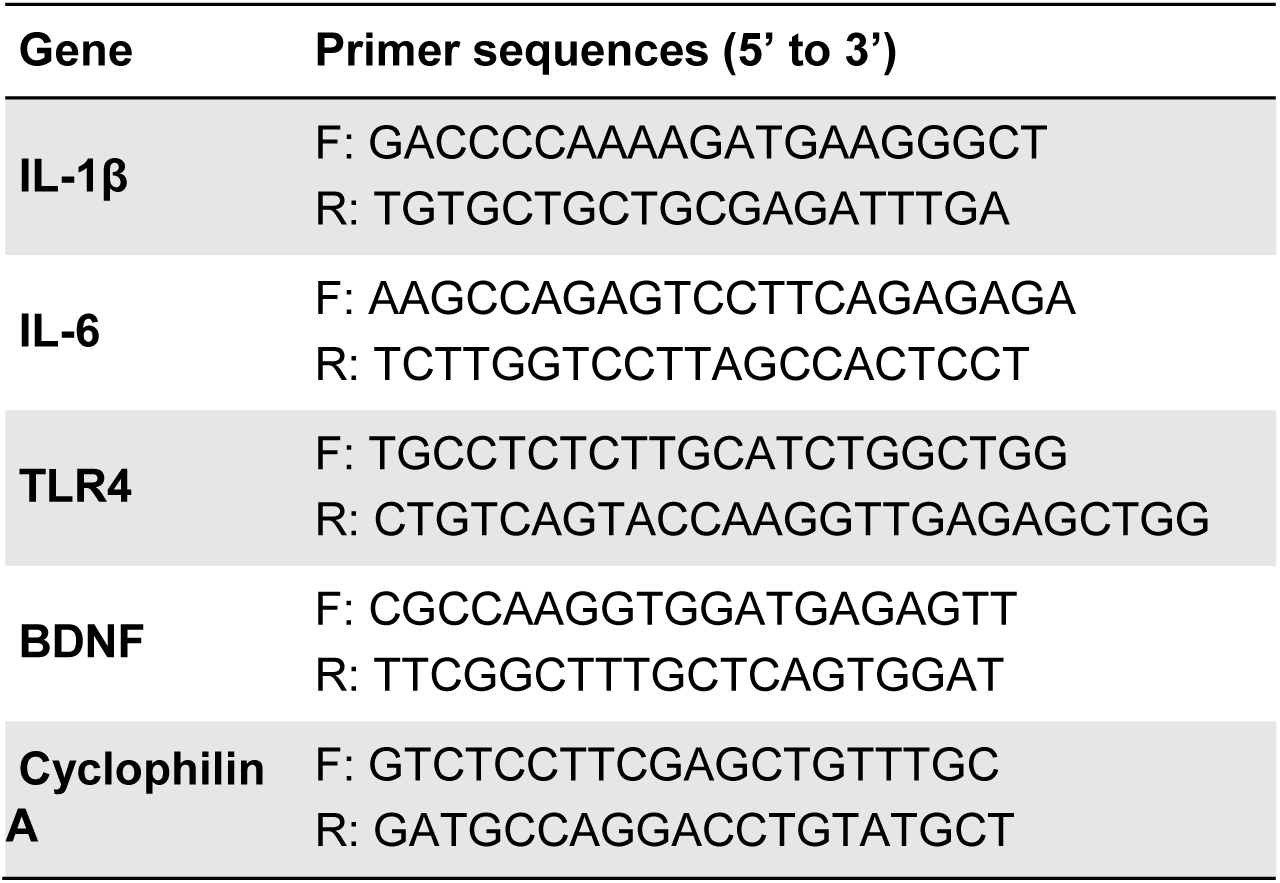
Nucleotide sequences of the primers used for the RT-PCR of genes.

## References

Best, L. M., Williams, B., Le Foll, B., Mansouri, E., Bazinet, R. P., Lin, L., De Luca, V., Lagzdins, D., Rusjan, P., Tyndale, R. F., Wilson, A. A., Hendershot, C. S., Heilig, M., Houle, S., Tong, J., Kish, S. J., & Boileau, I. (2020). Lower brain fatty acid amide hydrolase in treatment-seeking patients with alcohol use disorder: a positron emission tomography study with [C-11]CURB. Neuropsychopharmacology 2020 45:8, 45(8), 1289–1296. 10.1038/s41386-020-0606-2

Bilbao, A., Serrano, A., Cippitelli, A., Pavón, F. J., Giuffrida, A., Suárez, J., García-Marchena, N., Baixeras, E., Gómez de Heras, R., Orio, L., Alén, F., Ciccocioppo, R., Cravatt, B. F., Parsons, L. H., Piomelli, D., & Rodríguez de Fonseca, F. (2016). Role of the satiety factor oleoylethanolamide in alcoholism. Addiction Biology, 21(4), 859–872. 10.1111/ADB.12276/SUPPINFO

Carvalho, A. F., Heilig, M., Perez, A., Probst, C., & Rehm, J. (2019). Alcohol use disorders. The Lancet, 394(10200), 781–792. 10.1016/S0140-6736(19)31775-1

Cinar, R., Iyer, M. R., & Kunos, G. (2020). The therapeutic potential of second and third generation CB1R antagonists. Pharmacology & Therapeutics, 208, 107477. 10.1016/J.PHARMTHERA.2020.107477

Comerota, M. M., Gedam, M., Xiong, W., Jin, F., Deng, L., Wang, M. C., Wang, J., & Zheng, H. (2023). Oleoylethanolamide facilitates PPARα and TFEB signaling and attenuates Aβ pathology in a mouse model of Alzheimer’s disease. Molecular Neurodegeneration, 18(1), 56. 10.1186/S13024-023-00648-X/FIGURES/8

de Bruin, N. M. W. J., Lange, J. H. M., Kruse, C. G., Herremans, A. H., Schoffelmeer, A. N. M., van Drimmelen, M., & De Vries, T. J. (2011). SLV330, a cannabinoid CB1 receptor antagonist, attenuates ethanol and nicotine seeking and improves inhibitory response control in rats. Behavioural Brain Research, 217(2), 408–415. 10.1016/J.BBR.2010.11.013

Economidou, D., Mattioli, L., Cifani, C., Perfumi, M., Massi, M., Cuomo, V., Trabace, L., & Ciccocioppo, R. (2006). Effect of the cannabinoid CB1 receptor antagonist SR-141716A on ethanol self-administration and ethanol-seeking behaviour in rats. Psychopharmacology, 183(4), 394–403. 10.1007/S00213-005-0199-9/FIGURES/4

Garcia-Marchena, N., Pavon, F. J., Pastor, A., Araos, P., Pedraz, M., Romero-Sanchiz, P., Calado, M., Suarez, J., Castilla-Ortega, E., Orio, L., Boronat, A., Torrens, M., Rubio, G., de la Torre, R., Rodriguez de Fonseca, F., & Serrano, A. (2017). Plasma concentrations of oleoylethanolamide and other acylethanolamides are altered in alcohol-dependent patients: effect of length of abstinence. Addiction Biology, 22(5), 1366–1377. 10.1111/ADB.12408

González-Portilla, M., Moya, M., Montagud-Romero, S., de Fonseca, F. R., Orio, L., & Rodríguez-Arias, M. (2023). Oleoylethanolamide attenuates the stress-mediated potentiation of rewarding properties of cocaine associated with an increased TLR4 proinflammatory response. Progress in Neuro-Psychopharmacology and Biological Psychiatry, 124, 110722. 10.1016/J.PNPBP.2023.110722

Hansson, A. C., Gründer, G., Hirth, N., Noori, H. R., Spanagel, R., & Sommer, W. H. (2019). Dopamine and opioid systems adaptation in alcoholism revisited: Convergent evidence from positron emission tomography and postmortem studies. Neuroscience & Biobehavioral Reviews, 106, 141–164. 10.1016/J.NEUBIOREV.2018.09.010

Ho, W. S. V., Barrett, D. A., & Randall, M. D. (2008). ‘Entourage’ effects of N-palmitoylethanolamide and N-oleoylethanolamide on vasorelaxation to anandamide occur through TRPV1 receptors. British Journal of Pharmacology, 155(6), 837–846. 10.1038/BJP.2008.324

Hungud, B. L., Szakall, I., Adam, A., Basavarajappa, B. S., & Vadasz, C. (2003). Cannabinoid CB1 receptor knockout mice exhibit markedly reduced voluntary alcohol consumption and lack alcohol-induced dopamine release in the nucleus accumbens. Journal of Neurochemistry, 84(4), 698–704. 10.1046/J.1471-4159.2003.01576.X

Jin, P., Yu, H. L., Tian-Lan Zhang, F., & Quan, Z. S. (2015). Antidepressant-like effects of oleoylethanolamide in a mouse model of chronic unpredictable mild stress. Pharmacology Biochemistry and Behavior, 133, 146–154. 10.1016/J.PBB.2015.04.001

Kathryn Mchugh, R., & Weiss, R. D. (2019). Alcohol Use Disorder and Depressive Disorders. Alcohol Research : Current Reviews, 40(1), e1–e8. 10.35946/ARCR.V40.1.01

Lazenka, M. F., Selley, D. E., & Sim-Selley, L. J. (2013). Brain regional differences in CB1 receptor adaptation and regulation of transcription. Life Sciences, 92(8–9), 446–452. 10.1016/J.LFS.2012.08.023

Masana, Y., Wanaka, A., Kato, H., Asai, T., & Tohyama, M. (1993). Localization of trkB mRNA in postnatal brain development. Journal of Neuroscience Research, 35(5), 468–479. 10.1002/JNR.490350503

Moya, M., San Felipe, D., Ballesta, A., Alén, F., Rodríguez de Fonseca, F., García-Bueno, B., Marco, E. M., & Orio, L. (2021). Cerebellar and cortical TLR4 activation and behavioral impairments in Wernicke-Korsakoff Syndrome: Pharmacological effects of oleoylethanolamide. Progress in Neuro-Psychopharmacology and Biological Psychiatry, 108, 110190. 10.1016/J.PNPBP.2020.110190

Orio, L., Alen, F., Pavón, F. J., Serrano, A., & García-Bueno, B. (2019). Oleoylethanolamide, neuroinflammation, and alcohol abuse. Frontiers in Molecular Neuroscience, 11, 490. 10.3389/FNMOL.2018.00490/BIBTEX

Pagotto, U., Marsicano, G., Cota, D., Lutz, B., & Pasquali, R. (2006). The Emerging Role of the Endocannabinoid System in Endocrine Regulation and Energy Balance. Endocrine Reviews, 27(1), 73–100. 10.1210/ER.2005-0009

Peters, K. Z., Cheer, J. F., & Tonini, R. (2021). Modulating the Neuromodulators: Dopamine, Serotonin, and the Endocannabinoid System. Trends in Neurosciences, 44(6), 464–477. 10.1016/J.TINS.2021.02.001

Ray, L. A., Bujarski, S., Grodin, E., Hartwell, E., Green, R. J., Venegas, A., Lim, A. C., Gillis, A., & Miotto, K. (2018). State-of-the-art behavioral and pharmacological treatments for alcohol use disorder. 10.1080/00952990.2018.1528265, 45(2), 124–140. 10.1080/00952990.2018.1528265

Rivera, P., Fernández-Arjona, M. del M., Silva-Peña, D., Blanco, E., Vargas, A., López-Ávalos, M. D., Grondona, J. M., Serrano, A., Pavón, F. J., Rodríguez de Fonseca, F., & Suárez, J. (2018). Pharmacological blockade of fatty acid amide hydrolase (FAAH) by URB597 improves memory and changes the phenotype of hippocampal microglia despite ethanol exposure. Biochemical Pharmacology, 157, 244–257. 10.1016/J.BCP.2018.08.005

Rivera, P., Silva-Peña, D., Blanco, E., Vargas, A., Arrabal, S., Serrano, A., Pavón, F. J., Bindila, L., Lutz, B., Rodríguez de Fonseca, F., & Suárez, J. (2019). Oleoylethanolamide restores alcohol-induced inhibition of neuronal proliferation and microglial activity in striatum. Neuropharmacology, 146, 184–197. 10.1016/J.NEUROPHARM.2018.11.037

Rodríguez De Fonseca, F., Navarro, M., Gómez, R., Escuredo, L., Nava, F., Fu, J., Murillo-Rodríguez, E., Giuffrida, A., Loverme, J., Gaetani, S., Kathuria, S., Gall, C., & Piomelli, D. (2001). An anorexic lipid mediator regulated by feeding. Nature 2001 414:6860, 414(6860), 209–212. 10.1038/35102582

Rodríguez De Fonseca, F., Roberts, A. J., Bilbao, A., Koob, G. F., & Navarro, M. (1999). Cannabinoid receptor antagonist SR141716A decreases operant ethanol self administration in rats exposed to ethanol-vapor chambers. Zhongguo Yao Li Xue Bao = Acta Pharmacologica Sinica, 20(12), 1109–1114. https://europepmc.org/article/med/11189201

Sagheddu, C., Torres, L. H., Marcourakis, T., & Pistis, M. (2020). Endocannabinoid-Like Lipid Neuromodulators in the Regulation of Dopamine Signaling: Relevance for Drug Addiction. Frontiers in Synaptic Neuroscience, 12. 10.3389/FNSYN.2020.588660

Sánchez-Marín, L., Pavón-Morón, F. J., Rodríguez de Fonseca, F., & Serrano, A. (2022). Attenuation of oleoylethanolamide-induced reduction of alcohol consumption in adult rats exposed intermittently to alcohol during adolescence. Neuroscience Letters, 781, 136670. 10.1016/J.NEULET.2022.136670

Sayd, A., Antón, M., Alén, F., Caso, J. R., Pavón, J., Leza, J. C., de Fonseca, F. R., García-Bueno, B., & Orio, L. (2015). Systemic Administration of Oleoylethanolamide Protects from Neuroinflammation and Anhedonia Induced by LPS in Rats. International Journal of Neuropsychopharmacology, 18(6), 1–14. 10.1093/IJNP/PYU111

Schwartz, G. J., Fu, J., Astarita, G., Li, X., Gaetani, S., Campolongo, P., Cuomo, V., & Piomelli, D. (2008). The Lipid Messenger OEA Links Dietary Fat Intake to Satiety. Cell Metabolism, 8(4), 281–288. 10.1016/J.CMET.2008.08.005

Shahen-Zoabi, S., Smoum, R., Bingor, A., Grad, E., Nemirovski, A., Shekh-Ahmad, T., Mechoulam, R., & Yaka, R. (2023). N-oleoyl glycine and N-oleoyl alanine attenuate alcohol self-administration and preference in mice. Translational Psychiatry 2023 13:1, 13(1), 1–8. 10.1038/s41398-023-02574-4

Silva-Peña, D., Rivera, P., Alén, F., Vargas, A., Rubio, L., García-Marchena, N., Pavón, F. J., Serrano, A., De Fonseca, F. R., & Suárez, J. (2019). Oleoylethanolamide modulates BDNF-ERK signaling and neurogenesis in the hippocampi of rats exposed to δ9-THC and ethanol binge drinking during adolescence. Frontiers in Molecular Neuroscience, 12, 442287. 10.3389/FNMOL.2019.00096/BIBTEX

Soria-Gomez, E., Pagano Zottola, A. C., Mariani, Y., Desprez, T., Barresi, M., Bonilla-del Río, I., Muguruza, C., Le Bon-Jego, M., Julio-Kalajzić, F., Flynn, R., Terral, G., Fernández-Moncada, I., Robin, L. M., Oliveira da Cruz, J. F., Corinti, S., Amer, Y. O., Goncalves, J., Varilh, M., Cannich, A., … Bellocchio, L. (2021). Subcellular specificity of cannabinoid effects in striatonigral circuits. Neuron, 109(9), 1513–1526.e11. 10.1016/J.NEURON.2021.03.007

Yang, L. C., Guo, H., Zhou, H., Suo, D. Q., Li, W. J., Zhou, Y., Zhao, Y., Yang, W. S., & Jin, X. (2015). Chronic oleoylethanolamide treatment improves spatial cognitive deficits through enhancing hippocampal neurogenesis after transient focal cerebral ischemia. Biochemical Pharmacology, 94(4), 270–281. 10.1016/J.BCP.2015.02.012

Zhou, Y., Schwartz, B. I., Giza, J., Gross, S. S., Lee, F. S., & Kreek, M. J. (2017). Blockade of alcohol escalation and “relapse” drinking by pharmacological FAAH inhibition in male and female C57BL/6J mice. Psychopharmacology, 234(19), 2955–2970. 10.1007/S00213-017-4691-9/FIGURES/8

